# Quantification of tissue creatine content using capillary electrophoresis

**DOI:** 10.1101/2024.04.09.588648

**Authors:** Y. Jane Choi, Angelo P Bautista, Jessica R Terrill, J. Jane Pillow, Peter G Arthur

## Abstract

Creatine plays a fundamental role in cellular energy homeostasis. The current protocol describes an alternative method for creatine quantification in biological tissue samples using capillary electrophoresis, with high separation efficiency that enables rapid analysis with low sample volumes. The protocol involves homogenization of snap-frozen tissue in phosphate buffer, followed by electrophoresis through a bare-fused capillary (75 µm internal diameter) and measurement at 200 nm on the Agilent 7100 CE system. Under the optimised conditions, there was excellent linearity in creatine standards between 6.3 – 100 µM. The overall intra-assay variability for concentrations between 6.3 – 100 µM was 1.5 %, and the inter-assay variability was 6.4 %, with a limit of detection at 6 nmol/mg protein. The protocol was further benchmarked against a commercially available enzyme assay kit using lung samples from lambs that received continuous creatine or saline supplementation. There was good agreement between the two methods (mean difference = 0.42 [-0.26-1.1] nmol/mg protein). Importantly, capillary electrophoresis enables reliable detection of creatine in biological samples from just ∼1.5 mg of wet-weight lung tissue. Capillary electrophoresis enables rapid (<10 minutes) and highly efficient analysis of tissue samples and avoids challenges faced with traditional enzymatic assays. The current protocol was developed and optimised with ovine lung tissue, but it can be easily adapted to analyse various tissue types. For tissues with higher baseline creatine content, such as the skeletal muscles or brain, <1 mg wet weight tissue would be sufficient to detect creatine using capillary electrophoresis.

## Introduction

Creatine is a popular nutritional supplement with a fundamental role in energy homeostasis [1]. The creatine (Cr)/phosphocreatine (PCr) system is intrinsically involved in cellular energy homeostasis by converting PCr to Cr, which yields a high-energy phosphate group used to convert adenosine diphosphate (ADP) to adenosine triphosphate (ATP). Increasing the tissue creatine content, also called creatine ‘loading’, increases the intracellular pool of Cr and PCr and provides an alternate pathway to generate ATP rapidly [2]. In skeletal muscles, creatine exerts ergogenic effects in healthy individuals and therapeutic benefits in various myopathies [1,3]. In the brain, the benefit of creatine in neurodegenerative diseases (such as Huntington’s and Alzheimer’s disease) through improved cerebral bioenergetics is well-described [4]. More recently, there is growing evidence for the role of creatine as a perinatal neuroprotectant via energy preservation and antioxidant protection of the fetal brain during ischemic conditions [5,6].

Several methods are available to measure the creatine content of biological samples. Direct determination of creatine is often used in a clinical context to diagnose inborn errors of creatine metabolism [7]. These quantitative methods involve the analysis of patient serum or urine samples for creatine content via high-performance liquid chromatography [8] or mass-spectrometry-based approaches [9–13]. In a research context, similar HPLC methods are used to assess creatine content in tissue or cell culture samples [14–16] or ^31^P-magnetic resonance spectroscopy (MRS) to quantify high-energy phosphates in the brain tissue [17]. Alternatively, enzymatic detection of creatine involves fluorometric assay initially described by Passonneau and Lowry, which involves a multi-step enzymatic reaction to infer creatine concentration from the decrease in NADH fluorescence [18]. This method was used to determine creatine content in various tissue types, such as the muscle, brain, heart, liver and kidneys, from murine and ovine models [19–21]. However, enzymatic assays can be affected by the presence of interfering substances in the sample, and often require specific acid extraction during sample preparation that can be time-consuming.

We present an alternative method for creatine quantification in biological tissue samples using capillary electrophoresis. The advantage of capillary electrophoresis includes high separation efficiency that enables cost-effective and rapid analysis with low (nanolitre) sample volumes [22,23]. These features make capillary electrophoresis amenable to use with human biopsy samples. Capillary electrophoresis provides improved sensitivity at lower sample concentrations and avoids interference in enzymatic assays. While the detection of creatine and creatinine using CE was described previously [24], existing methods are limited to human urinary creatine and creatinine levels. The current protocol describes workflow for determination of creatine content from snap-frozen tissue samples; from sample homogenisation and extraction to optimised conditions for capillary electrophoresis. We developed and optimised the current protocol with ovine lung samples to determine changes in tissue creatine content after a direct fetal creatine supplementation [25]. However, the protocol can be easily adapted to analyse various tissue types.

## Materials and Methods

The protocol described in this peer-reviewed article is published on protocols.io (dx.doi.org/10.17504/protocols.io.4r3l22j23l1y/v2) and is included for printing as Supporting Information file 1 with this article.

The creatine peaks on the electropherogram were identified via enzymatic depletion of the signal and enhancement of the endogenous signal after spiking with additional creatine. Creatine concentration was calculated as the area under the peak (AUC) of the electropherogram and expressed as a ratio compared to the reference DMSO signal. Standards were injected in technical duplicates and run at the beginning of each day before sample analysis. The standards were repeated across four days, and a standard curve was generated from each day to assess the inter-assay variability. Samples were standardised to 1 mg/mL protein concentration and injected in technical duplicates to assess the intra-assay variability. Creatine concentration was interpolated from the standard curve and expressed as nmol/mg protein. The results from capillary electrophoresis were subsequently benchmarked against a commercially available creatine assay kit (MAK079, Sigma-Aldrich), which measures creatine without phosphocreatine due to a lack of creatine phosphokinase. Samples were similarly standardised to 1 mg/mL, and the assay was performed according to the manufacturer’s guidelines. Creatine concentrations are similarly expressed as nmol/mg protein.

### Expected results

Using optimised conditions described in the protocol, creatine peak is confirmed after depletion with creatinase enzyme and increased after spiking with additional creatine (**Fig 1**).

**Fig 1.**
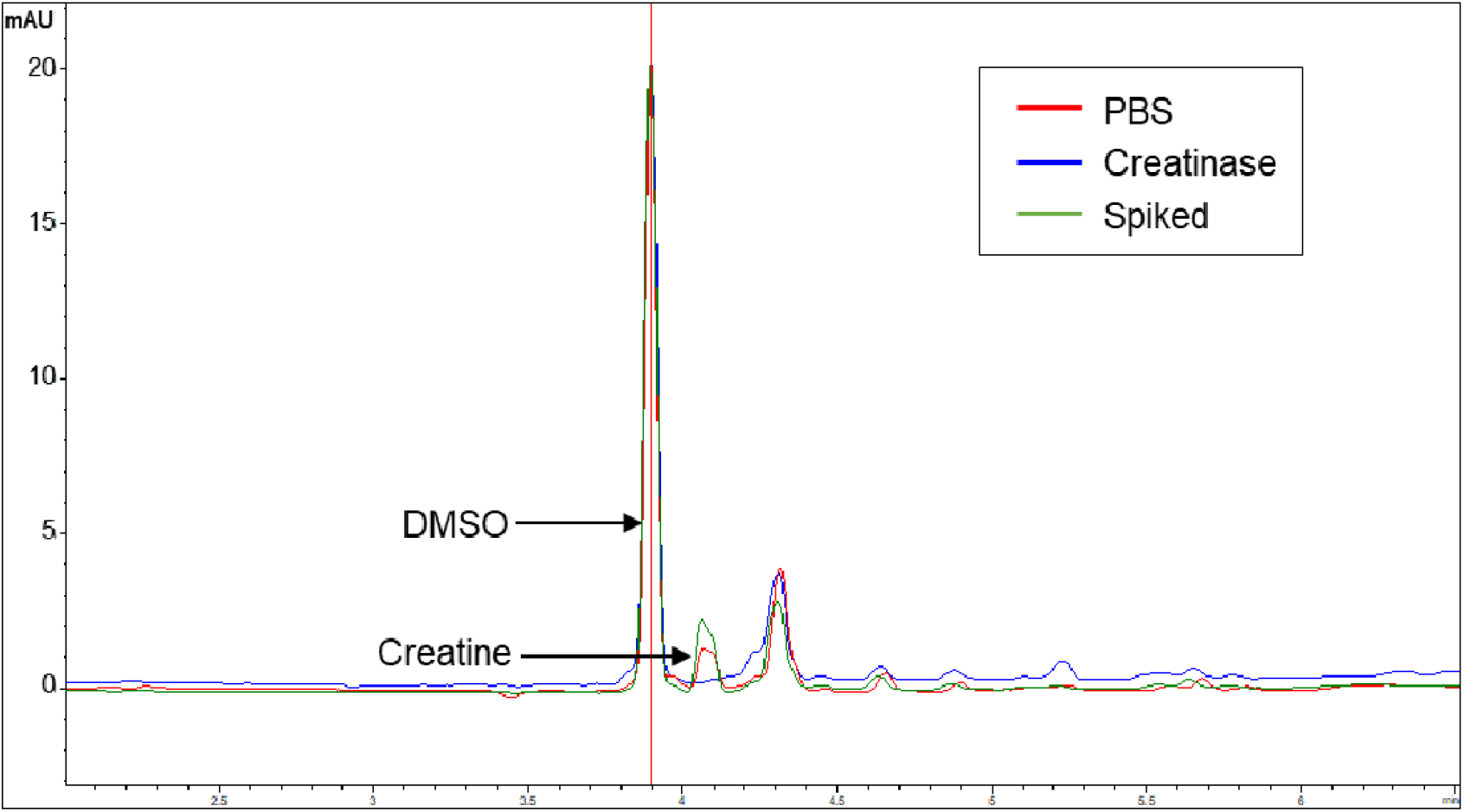
Identification of creatine peak by capillary electrophoresis under optimised conditions with UV detection at 200 nm. Snap-frozen lung tissues were homogenised in PBS with 0.01 % DMSO (red, baseline), and the creatine peak on the electropherogram was identified via enzymatic depletion with creatinase (blue) and spiking (green) with additional creatine (25 µM). A bare-fused capillary (75 µm internal diameter) at an effective length of 36.5 cm (length to detector 8.5 cm) was used. The background electrolyte contained 75 mM NaH_2_PO_4_ and 150 mM SDS at pH 6. A voltage of 15 kV was applied at 25 °C with sample injection at 10 mbar for 10 s.

### Linearity and assay variability

We assessed the separation performance via the linearity of the standard curve and the intra- and inter-assay variability. The concentration range for the assessment of linearity was determined based on the expected range of concentration for this sample type. A representative image of the electropherograms generated at each standard concentration is illustrated in **Fig 2**, and the linear regression parameters are described in **Fig 3**. Under the optimised conditions tested, there was excellent linearity in standards between 6.3 – 100 µM.

**Fig 2.**
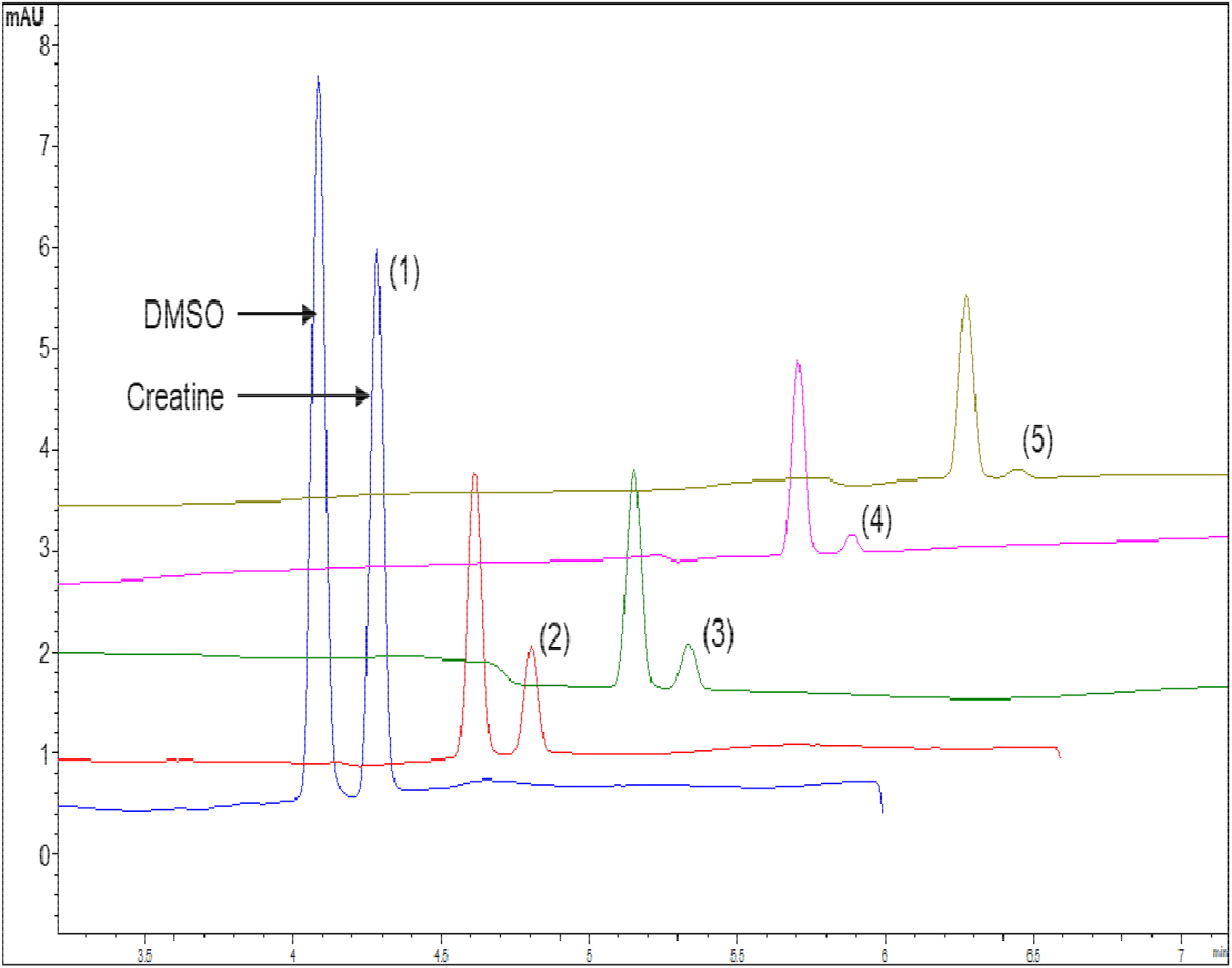
Representative electropherograms of creatine standards under optimised CE conditions with UV detection at 200 nm. The creatine peaks are identified as: (1) 100 µM, (2) 50 µM, (3) 25 µM, (4) 12.5 µM, (5) 6.3 µM, (6) 0 µM. Creatine standards were prepared in homogenisation buffer, which contains 50 mM NaH_2_PO_4_, 150 mM SDS and 0.01 % DMSO (as the reference signal) at pH 6. The separation conditions were identical to Fig 1.

**Fig 3.**
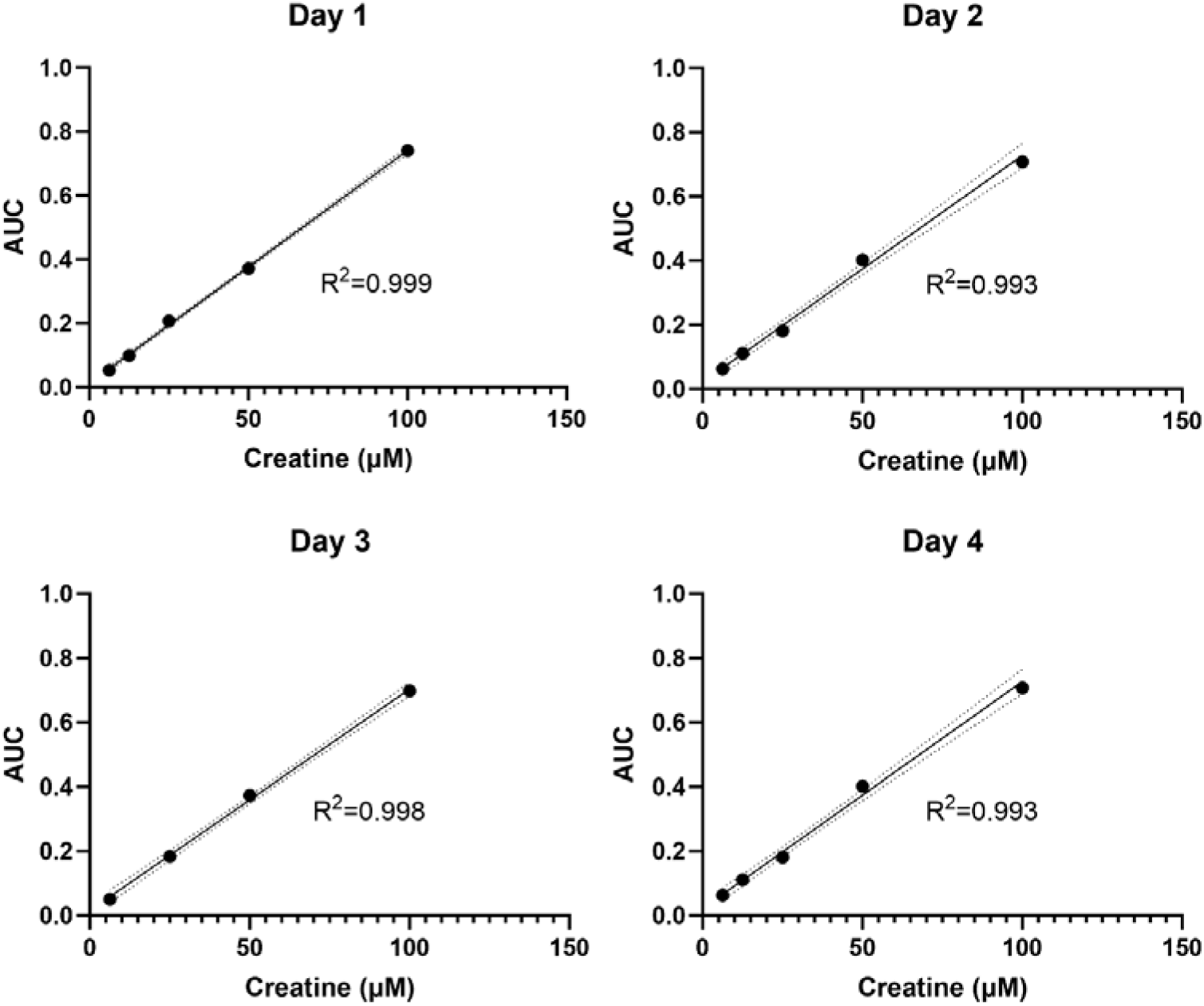
Linearity of creatine standards between 6.3 – 100 µM. Each panel illustrates the standard curve generated on four separate days. The dotted line represents the 95 % confidence interval of the linear regression. The separation conditions were identical to Fig 1.

The overall intra-assay variability for concentrations between 6.3 – 100 µM was 1.5 %, and the inter-assay variability was 6.4 %. The limit of detection was 6 nmol/mg protein, but variables such as the concentration of the sample injected or the capillary diameter could be changed to enhance the lower limit of detection.

### Determination of creatine content in fetal lamb lungs and benchmarking against commercial enzymatic assay kit

The optimised conditions were used to compare creatine content in ovine fetal lungs in groups that received a direct intravenous supplementation of creatine (6 mg kg^-1^hr^-1^) or saline (0.9 % heparinized saline [2 IU/mL]) for 17 days before surgical delivery at 110 days gestation [25]. We tested a random subset of animals from both saline control and creatine-supplemented groups (n = 5/group, based on *a priori* calculation). All samples had creatine levels within the range of detection. Creatine supplementation significantly increased creatine content in the fetal lungs, determined using a commercial enzymatic assay (mean difference = 11.3 [8.9, 13.7] nmol/mg protein; p < 0.0001), and capillary electrophoresis (mean difference [95 % CI] = 11.8 [8.7, 14.9] nmol/mg protein; p < 0.0001) (**Fig 4A**). Fig 4B illustrates a Bland-Altman comparison of the two methods. The two methods showed good agreement, although capillary electrophoresis derived a marginally higher creatine content compared to the enzymatic assay (mean difference [95% CI] = 0.42 [-0.26 – 1.1] nmol/mg protein). While this bias was not statistically significant, it could be explained by potential interference in enzyme assay or lack of completion of the reaction [18]. Regardless, the mean difference was within the pre-determined range of ± 15 %, and both methods showed a similar magnitude of difference between groups that did and did not receive creatine supplementation.

**Fig 4.**
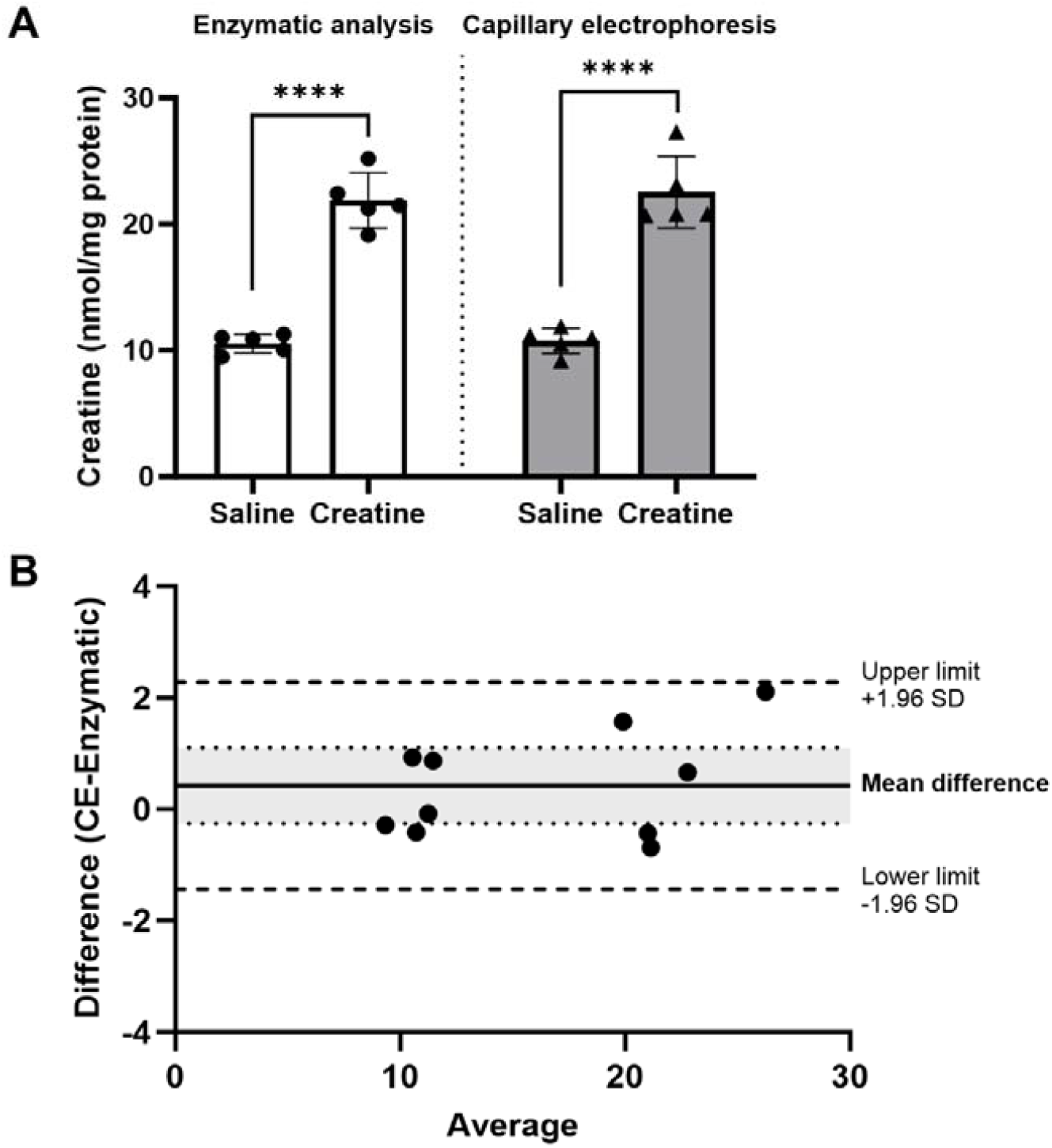
Comparison of capillary electrophoresis (CE) and commercial enzymatic assay kit. (A) Creatine content in the lung tissue was determined using a commercial enzymatic assay kit (MAK079, Sigma-Aldrich) and capillary electrophoresis under separation conditions described in Fig 1. Groups were compared using unpaired t-tests and presented as mean ± SD. ****p<0.0001. (B) A Bland-Altman comparison between two methods. The dotted lines on either side of the mean difference denote the 95 % confidence interval of the mean difference.

### Comments on the utility of this protocol

Overall, the current protocol provides a streamlined process to analyse creatine content in biological tissue samples. Given the excellent sensitivity of the protocol at low tissue creatine concentrations, we calculated that approximately 1.5 mg tissue (wet weight) would be sufficient to determine creatine content in a lung sample (assuming ∼10 nmol/mg protein in control conditions and that each mg of tissue yields ∼0.2 mg protein). The current protocol was developed and optimised with ovine lung tissue samples, but it can be easily adapted to analyse various tissue types. We anticipate that even less amount of tissue would be sufficient (< 1 mg) to reliably detect creatine in tissues with higher baseline creatine content, such as the skeletal muscles or brain.

## Ethics declarations

Animal ethics approval for the collection of fetal lung tissue was obtained at the University of Western Australia (RA/3/100/1607), and animal research was conducted in accordance with the National Health and Medical Research Council Code of Practice for the Care and Use of Animals for Scientific Purposes (Eighth Edition).

## Supporting information

S1: Step-by-step protocol, also available on protocols.io.

## Acknowledgements

The schematic illustration of the protocol workflow was created with Biorender.com.

## Authors contributions

YJC conceptualised, performed the experiments, drafted the protocol and the manuscript. AB conceptualised, performed the experiments and reviewed the protocol. JT interpreted results and reviewed the protocol. JJP obtained funding for the study and reviewed the protocol. PGA conceptualised, interpreted the results, and reviewed the protocol. All authors reviewed and revised the manuscript and approved the final form.

