## Supporting Information File 1 for "Quantification of tissue creatine content using capillary electrophoresis" (from another version of this preprint: version 1, 2024-04-10)

DBVU2N6W

**Protocol Info:** Jane Choi, Angelo Bautista, Jessica Terrill, J. Jane Pillow, Peter G Arthur . Quantification of tissue creatine content using capillary electrophoresis. **protocols.io** https://protocols.io/view/quantification-of-tissue-creatine-content-using-ca-dbvu2n6wVersion created by Jane Choi

**Created:** Apr 09, 2024

**Last Modified:** Apr 09, 2024

**PROTOCOL integer ID:** 97940

**Keywords:** creatine, capillary electrophoresis, lung, skeletal muscle, biopsy

**Funders Acknowledgement:**
Australian National Health & Medical Research Council
Grant ID: GNT1196188
Stan Perron Foundation
Grant ID: Post-PhD Career Launching Award

### Quantification of tissue creatine content using capillary electrophoresis V.(dbvu2n6w)

Jane Choi[1,2], Angelo Bautista[1], Jessica Terrill[1], J. Jane Pillow[1,2], Peter G Arthur[1]

[1]University of Western Australia; [2]Telethon Kids Institute

Jane Choi
Jane Choi
University of Western Australia

#### DISCLAIMER

No competing interests are declared.

#### ABSTRACT

The current protocol describes an alternative method for creatine quantification in biological tissue samples using capillary electrophoresis, with high separation efficiency that enables rapid analysis (<10 minutes) with low sample volumes. The protocol involves homogenization of snap-frozen tissue samples in phosphate buffer, followed by electrophoresis through a bare-fused capillary (75 µm i.d.) and measurement at 200 nm. Under the optimised conditions, we found excellent linearity in creatine standards between 6.3 – 100 µM. The overall intra-assay variability for concentrations between 6.3 – 100 µM was 1.5 %, and the inter-assay variability was 6.4 %, with a limit of detection at 6 nmol/mg protein. Variables such as the amount of sample injected or the capillary diameter could be changed to enhance the lower limit of detection. The current protocol was developed and optimised using ovine lung tissue samples, but it can be easily adapted to analyse various tissue types.

#### PROTOCOL REFERENCES

Adams DS. Making solutions from hydrated compounds. Cold Spring Harbor Protocols. 2008 May 1;2008(5):pdb-ip54.

#### GUIDELINES

This protocol describes the capillary electrophoresis method optimised for detecting creatine from tissue samples. The current parameters are optimised for ovine snap-frozen lung parenchyma. However, the method can be further optimised for various different tissue types (validation in progress).

MATERIALS

**Equipment**

- Agilent 7100 Capillary Electrophoresis machine
- Agilent cassette #G7100-60002
- Agilent vial 250 uL#5190-3155
- Agilent polyurethane snap caps, #5042-6491
- Bare-fused silica capillary (75 µm inner diameter; Polymicro Technologies, AZ, USA)
- Tissue homogeniser - TissueRuptor or TissueLyser

**Chemicals**

- Sodium phosphate monobasic ($NaH_2PO_4$)
- Sodium dodecyl sulfate (SDS)
- Dimethylsulfoxide (DMSO)
- NaOH pellets
- PBS tablets
- Creatinase, Merck #C2409
- Creatine kinase, Roche #10127566001
- Creatine monohydrate, Merck #C3630
- Creatine anhydrous, Merck #C0780

#### Preparation of buffers

1 Prepare the stock phosphate buffer (PB) [M] 200 millimolar (mM) PB pH 6

1.1 Add 1199.8 mg $NaH_2PO_4$ to 30 mL dd$H_2O$, dissolve well

1.2 pH the solution to 6.0 with concentrated NaOH

1.3 top up the volume to 50 mL with dd$H_2O$ (ideally using a volumetric flask)

1.4 filter sterilise the solution through a 0.45 µm syringe filter

1.5 store at 4 °C for a maximum of 4 weeks. Discard if any precipitate forms.

2 M × V × #$H_2O$ × 18.015 = mL $H_2O$ contributed by the hydrated compound.

3 Prepare the background electrolyte (BGE) [M] 75 millimolar (mM) PB [M] 150 millimolar (mM) SDS pH 6

3.1 Dilute 18.75 mL of 200 mM PB buffer (pH = 6) with 20 mL dd$H_2O$

3.2 Add 2.163 g SDS to the solution and dissolve well

3.3 Top up the volume to 50 mL with $ddH_2O$ using a volumetric flask

3.4 Filter sterilise the solution through a 0.45 µm syringe filter

3.5 Store at Room temperature for up to a week. Discard if any precipitate forms.

4 Prepare the homogenisation buffer (S-BGE) [M] 50 millimolar (mM) PB [M] 150 millimolar (mM) SDS [M] 0.01 % volume DMSO pH 6

4.1 Dilute 12.5 mL of 200 mM PB buffer (pH=6) with 17.5 mL $ddH_2O$

4.2 Add 2.163 g SDS to the solution and dissolve well

4.3 Top up the volume to 50 mL with $ddH_2O$ using a volumetric flask

4.4 Remove 5 µL of solution and replace with 5 µL DMSO

4.5 Filter sterilise the solution through a 0.45 µm syringe filter

4.6 Store at Room temperature for up to a week. Discard if any precipitate forms.

#### Preparation of creatine standard

5 Prepare creatine standards by dissolving creatine monohydrate (C3630, Merck) or anhydrous (C0780, Merck) to a stock concentration of 10 mM.
*Note*: If using creatine monohydrate – account for the hydrated compound by excluding the mass of $H_2O$ from calculations (Adams, 2008). Although this is ***negligible*** at small volumes/concentrations.

$$M \times V \times \#H_2O \times 18.015 = mL\ H_2O \text{ contributed by the hydrated compound.}$$

5.1 Dilute the 10 mM stock solution to the appropriate range using the **same solvent as the samples**. i.e., If running enzymatic analysis – dilute the standards in PBS. If running samples – dilute the standards in the S-BGE (50 mM PB buffer).
Note: Prepare creatine standards within the range typically expected for the tissue type. However, this can also depend on the protein concentration of the samples.
For lung tissue, standard curve concentrations between the 6.25 - 100 µM range have shown excellent linearity with low inter-assay variability.

#### Sample preparation for enzymatic analysis

6 This step is required once for each sample type to determine and confirm which peak on the electropherogram corresponds to creatine. Validation is performed by adding creatinase enzyme (which breaks down creatine into urea & sarcosine) to deplete the signal. Comparison can be made between samples with and without creatinine to identify the modified peak as creatine.
In essence, the addition of the creatinase enzyme will essentially serve as a negative control.

6.1 Rapidly homogenise snap-frozen tissue samples in 20 volume of 1X PBS
  a. With a mortar & pestle with liquid nitrogen
  b. Or a bead beater

6.2 Centrifuge the homogenates 10000 x g, 4°C, 00:05:00 5m

6.3 Transfer supernatants into a new tube.

6.4 Take an aliquot of the supernatant and determine the protein concentration with an appropriate protein assay (e.g. BCA, Bradford, DC assay)

6.5 Dilute the samples with PBS to standardise the protein concentration in all samples (typically 2 mg/mL for lungs).

6.6 Aliquot and store at -80 °C until use.

#### Sample preparation for CE analysis

7 Freshly prepared S-BGE is recommended for sample homogenisation

7.1 Rapidly homogenise snap-frozen tissue samples in 20 volume of S-BGE
  a. With a mortar & pestle with liquid nitrogen
  b. Or a bead beater

7.2 Centrifuge the homogenates 10000 x g, Room temperature, 00:05:00
*Note, keep samples in RT as SDS will precipitate at 4 degrees.*

7.3 Transfer supernatants into a new tube

7.4 Take an aliquot of the supernatant and determine the protein concentration with an appropriate protein assay (e.g. BCA, DC assay are compatible with SDS in the buffer)

7.5 Dilute the samples with S-BGE to standardise the protein concentration in all samples (typically 2 mg/mL for lungs).

7.6 Aliquot and store at -80 °C until use.

7.7

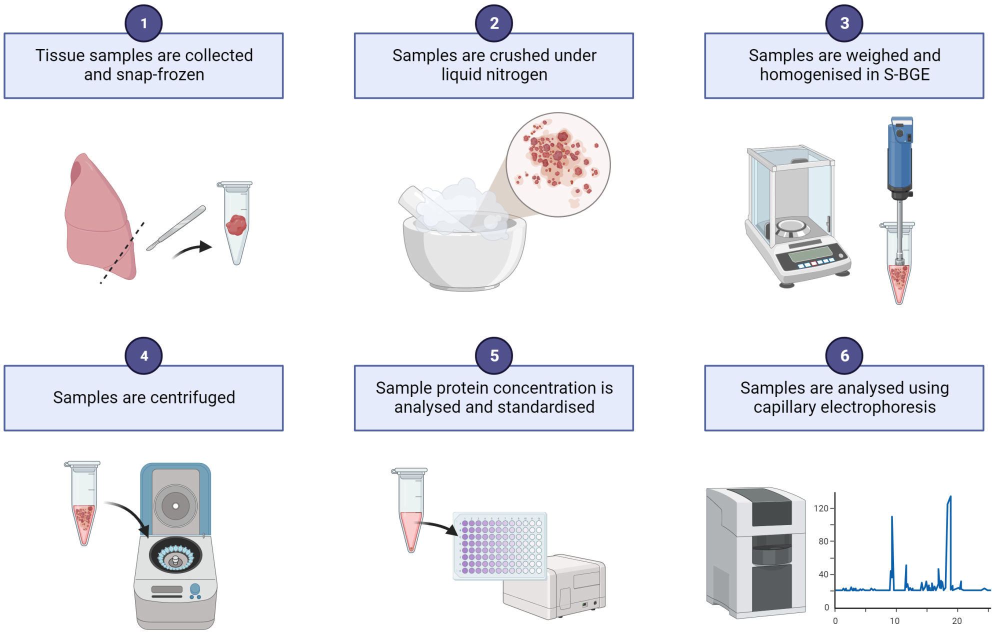


#### Equipment setup

8 Capillary electrophoresis is performed on the **Agilent 7100 CE system** (Agilent Technologies, CA, USA) with:

- bare-fused silica capillary (75 µm inner diameter; Polymicro Technologies, AZ, USA)
- effective length 36.5 cm
- length to detector 8.5 cm

*Note: wrap the capillary around the cassette so there is no overlap or touching, as this can generate and spread heat.*

9 Create a new protocol with the separation parameters as follows:

- Injection – 10 mbar for 10 s
- Separation is achieved with a constant voltage of 15 kV applied for 6 mins
- Capillary temperature set at 25 °C
- UV absorbance is measured at 200 nm

Post-conditioning parameters (after each sample) are:

1. dd$H_2O$ for 30 s
2. 0.1 M NaOH for 30 s
3. dd$H_2O$ for 30 s
4. BGE for 120 s

#### Running the Assay

10 At the beginning of each run, pre-condition the capillary by flushing in the following order:

1. $ddH_2O$ for 180 s
2. 0.1 M NaOH for 500 s
3. $ddH_2O$ for 120 s
4. BGE for 600 s

11 Load each tube with at least 10 µL (preferably 20 µL) of sample. Avoid creating bubbles as this can interfere with sample injection. Load the tubes on the Agilent machine.

12 Run the protocol. The samples can be submitted individually or as a sequence. If running more than ten samples, insert a flush protocol ($ddH_2O$ or BGE for 120 s) every ten runs.

Depending on the number of samples to analyse, it may be best to prepare two protocols with two different BGE & waste tubes assigned to ensure sufficient background buffer for the analysis and that the waste doesn't overflow.

13 At the end of the day, ensure post-conditioning has been successfully run and turn off the equipment.
